# A fully automated deep learning pipeline for high-throughput colony segmentation and classification

**DOI:** 10.1101/801845

**Authors:** Sarah H. Carl, Lea Duempelmann, Yukiko Shimada, Marc Bühler

## Abstract

**Background:** Adenine auxotrophy is a commonly used non-selective genetic marker in yeast research. It allows investigators to easily visualize and quantify various genetic and epigenetic events by simply reading out colony color. However, manual counting of large numbers of colonies is extremely time-consuming, difficult to reproduce and possibly inaccurate.

**Results:** Using cutting-edge neural networks, we have developed a fully automated pipeline for colony segmentation and classification, which speeds up white/red colony quantification 100-fold over manual counting by an experienced researcher.

**Conclusions:** Our approach uses readily available training data and can be smoothly integrated into existing protocols, vastly speeding up screening assays and increasing the statistical power of experiments that employ adenine auxotrophy.

## Background

Auxotrophy is the inability of an organism to synthesize a particular organic compound required for its growth. For example, yeasts with mutations in the adenine biosynthetic pathway cannot grow on media lacking adenine. When grown under adenine-limiting conditions, adenine auxotrophs grow but accumulate a cell-limited red pigment in their vacuoles, whereas wild type cells grow white. This red/white colony color assay has been widely used over the last decades for the investigation of many biological processes, such as recombination, copy number, chromosome loss, plasmid stability, prion propagation, or epigenetic gene regulation in both budding and fission yeasts. However, adapting this assay to quantitative high-throughput applications has proven challenging, as it requires extensive scoring of colony phenotypes by eye. In addition to being time-consuming and tedious, manual colony scoring may suffer from inaccuracy and irreproducibility. Nonetheless, up to now manual scoring is a common practice in the yeast community. Modern machine-learning techniques such as deep learning have made huge strides in automated image classification in recent years and are beginning to be applied to previously intractable problems in the biomedical imaging domain. We set out to leverage these recent developments to build a computational pipeline that would enable fully automated high-throughput adenine auxotrophy-based screening and quantification.

Typically, red/white colony color assays start with plating individual yeast cells on limiting adenine indicator plates and allowing them to grow until they form colonies large enough to be inspected by eye. Each plate may represent an independent condition or genotype, the penetrance of which can be assessed by quantifying the percentage of non-white colonies per plate. In order to create a pipeline that would fit into existing protocols as seamlessly as possible, we considered our input to be images of plates and our desired output to be percentages of white and non-white colonies per plate. Two major tasks are required to generate this output: separating the colonies from the plate background and classifying them individually as white or non-white.

These two tasks could conceivably be completed either in one step, as with a single-shot detector^1^ or RetinaNet^2^, or in two separate steps, such as with a semantic segmentation, where each pixel is assigned a label such as “foreground” or “background”, followed by classification of cropped images. While a single-step approach may be preferable from the perspective of algorithmic efficiency and speed^3^, the training data annotations are more complex, requiring both manually assigned labels and matched bounding boxes identifying the location of each colony on a plate. As insufficient training data is a common problem hampering efforts to apply deep learning in many biological domains^4^, we opted to use a pragmatic approach, treating the segmentation and classification steps as separate problems (Fig. 1a). This allowed us to use simpler and, when available, pre-existing annotations for training data: for the segmentation task, we used masks generated previously using the Ilastik image-processing toolkit^5^, while for the classification task, we relied on manual labels assigned by experienced biologists to cropped images of single colonies. All of our training data consisted of *Schizosaccharomyces pombe* colonies, with red pigment resulting from heterochromatin-mediated silencing of the *ade6*^*+*^ gene.

**Figure 1.**
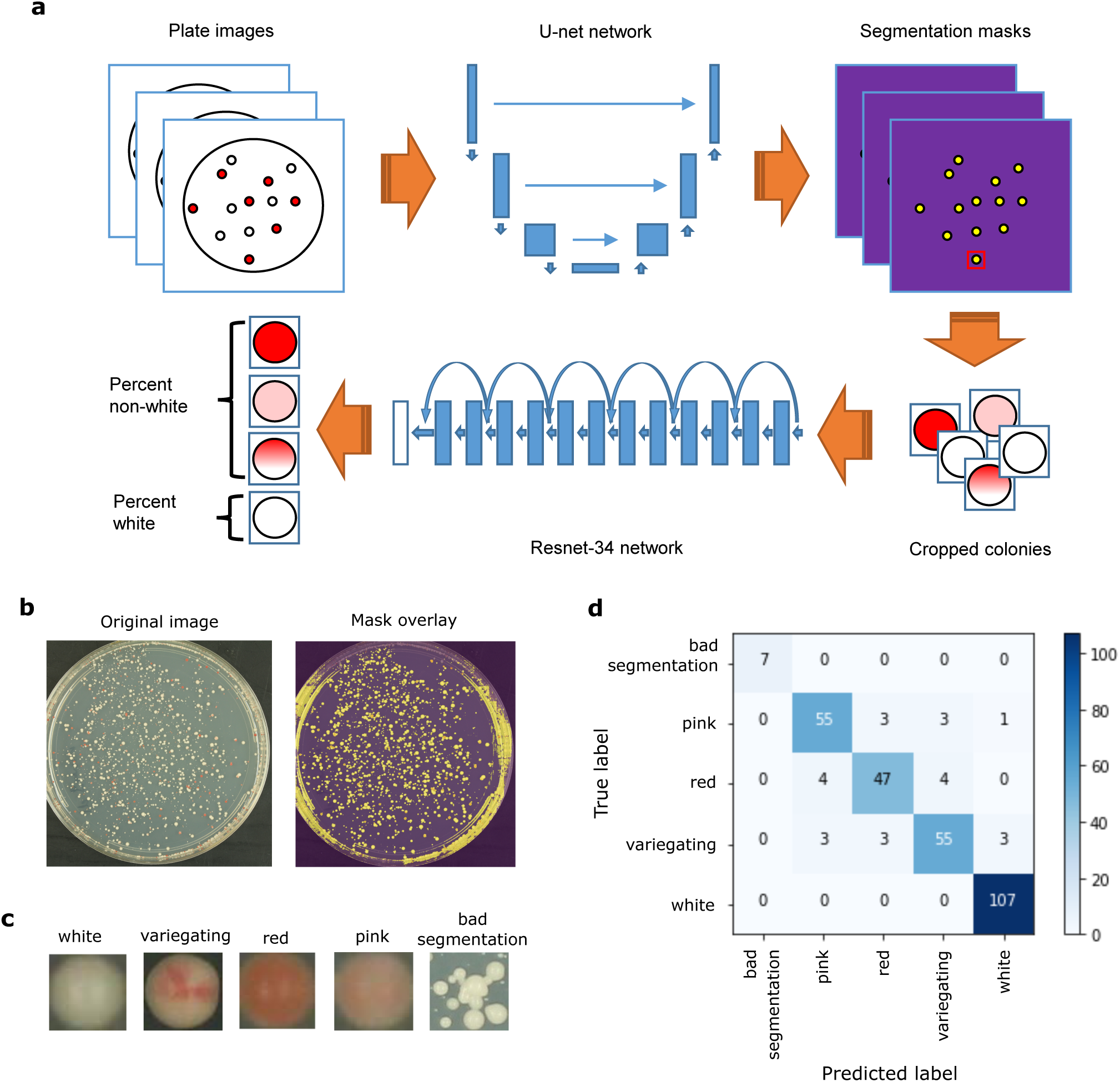
Overview of deep learning setup and results. a) Schematic of the entire automated colony classification pipeline. Plate images are given as input, then pass through a U-net network for segmentation prediction, resulting in cropped colony images. These are then fed into a Resnet-34 network for classification, followed by plate-level aggregation into white and non-white percentages. b) An example input plate image (left), and the same image overlaid with the predicted segmentation mask (right). c) Examples of cropped colonies classified into each of the 5 possible classes. d) Confusion matrix showing the results of the classification step (Resnet-34 network) on the validation data. Numbers in each square indicate the number of colonies with each true (y-axis) and predicted (x-axis) label.

### Implementation

All analyses were performed in Python v. 3.6.5 inside a conda virtual environment (v. 4.5.4). Deep learning models were built and trained using the fast.ai library (https://github.com/fastai/fastai)^6^. Image processing steps were performed using the scikit-image library (v. 0.13.1) (https://scikit-image.org/)^7^. Model training and prediction was run on a GeForce GTX 1080 GPU with CUDA v. 9.0 and CUDNN v. 7.1.2.

#### Image segmentation

A modified U-net architecture using a Resnet-34 network pre-trained on ImageNet as the encoder was used as the network architecture for the plate image segmentation task^8,9^. A total of 492 pairs of plate images and corresponding segmentation masks generated by Ilastik were used as training data, with approximately 20% (95 pairs) set aside for validation. Binary cross-entropy with logits (“BCEWithLogitsLoss”) was used as a loss function, and the dice coefficient was used as an evaluation metric. Data augmentation was applied to the training images with the following transformations: random rotation up to 4 degrees, random horizontal flip, random adjustment of brightness and contrast. The same transformations, except for adjustment of brightness and contrast, were applied to the training masks.

The network was first trained with center-cropped masks and images resized to 512×512 pixels, with a batch size of 4. A weight decay parameter of 1e-7 was used for all training. First, only the last layer was trained using stochastic gradient descent with restarts (SGDR) for 1 cycle of 8 epochs, using a circular learning rate scheduler with a maximum learning rate of 4e-2, a minimum learning rate of 8e-3, 1 epoch of increasing learning rate, and 7 epochs of decreasing learning rate^10,11^. Next, all layers were trained using SGDR for 1 cycle of 20 epochs. Differential learning rates were applied across layers, with the first 1/3 of layers having a maximum learning rate of 1e-4, the middle 1/3 having a maximum learning rate of 1e-3, and the last 1/3 having a maximum learning rate of 1e-2. A circular learning rate scheduler was again used, with minimum learning rates of 1/20^th^ of respective maximum learning rates, 2 epochs of increasing learning rates, and 18 epochs of decreasing learning rates. The resulting weights were then saved and used as a starting point to train the network with larger, 1024×1024 images.

Training images and masks were scaled up to 1024×1024 pixels and the network was further trained. First, only the last layer was trained using SGDR for 1 cycle of 2 epochs, using a circular learning rate scheduler with a maximum learning rate of 4e-2, a minimum learning rate of 8e-3, 0.5 epochs of increasing learning rate, and 1.5 epochs of decreasing learning rate. Next, all layers were trained using SGDR for 20 cycles of 20 epochs. Differential learning rates were applied across layers, with the first 1/3 of layers having a maximum learning rate of 4e-5, the middle 1/3 having a maximum learning rate of 2e-4, and the last 1/3 having a maximum learning rate of 4e-3. A circular learning rate scheduler was again used, with minimum learning rates of 1/20^th^ of respective maximum learning rates, 2.5 epochs of increasing learning rates per cycle, and 17.5 epochs of decreasing learning rates per cycle. The resulting weights were saved and used for prediction.

#### Colony classification

A Resnet-34 network that had been pre-trained on ImageNet was used as the network architecture for the colony classification task. The final output layer was replaced with a layer predicting 5 classes (“white”, “red”, “pink”, “variegating” and “bad segmentation”). A total of 1476 manually labeled, cropped images of individual colonies were used as training data, with approximately 20% (295 images) set aside for validation. Out of the total pool of images, 537 were labeled as “white”, 273 as “red”, 310 as “pink”, 318 as “variegating”, and 38 as “bad segmentation”. Validation images were chosen so as to have the same proportions among the five classes as in the total pool. For training, images were resized to 50×50 pixels and a batch size of 128 was used. Stochastic gradient descent (SGD) with a Momentum of 0.9 was used as an optimization algorithm. Categorical cross-entropy was used as a loss function, and both log loss and accuracy were used as evaluation metrics. Data augmentation was applied to the training images with the following transformations: random rotation up to 10 degrees, random rotation or reflection of dihedral group 8, random adjustment of brightness and contrast, random zoom up to 1.1x.

The last layer of the network was first trained without data augmentation for 1 epoch using a learning rate of 1e-2. Data augmentation was then added, and it was trained with SGDR for 3 cycles of 1 epoch each using cosine annealing, with an initial learning rate of 1e-2. Next, all layers were trained for 17 sets of 3 cycles of increasing length (1 epoch, followed by 2 epochs, followed by 4 epochs), for a total of 51 cycles and 119 epochs. Differential learning rates were applied across layers, with the first 1/3 of layers having a maximum learning rate of 1.1e-4, the middle 1/3 having a maximum learning rate of 3.3e-4, and the last 1/3 having a maximum learning rate of 1e-3. Training was stopped when over-fitting was observed, and the resulting weights were saved and used for prediction.

#### Mask prediction, post-processing and colony class prediction

The full prediction pipeline is implemented as follows: first, plate images are center-cropped and resized to 1024×1024 pixels. Segmentation masks are predicted using the trained U-net network, then resized to match the dimensions of the original (center-cropped) images. Border clearing and morphological opening are applied to the masks, reducing plate edge artefacts. Individual colonies are then labelled and a bounding box is drawn around each one, defining a region. Finally, colonies are selected if they have a regional eccentricity <= 0.6 and a regional area >= 400 pixels. Selected colonies are cropped and saved as individual .jpg images, retaining information about which plate each colony came from.

Classes are then predicted for individual colonies using the trained Resnet-34 network with test-time augmentation. This applies the same data augmentation transforms as were used during training to the test images, creating four randomly transformed versions of each image, then takes the average prediction for all four plus the original. The five colony class predictions are then aggregated per plate, and the percentage of non-white colonies is defined as the sum of predicted red, pink and variegating colonies divided by the sum of all properly segmented colonies (excluding those in the “bad segmentation” class). Segmentation and colony class prediction can also be performed separately, allowing for classification of previously-segmented images.

## Results

To perform the segmentation task, we chose a U-net-like architecture implemented in the fast.ai library^6^. U-net was developed specifically for semantic segmentation and has been successfully applied to complex biomedical images such as electron microscopy of neuronal structures and MRI or ultrasound images in breast cancer screening^9,12,13^. We trained the U-net network using 492 plate images and corresponding Ilastik-generated masks, with 20% kept aside for validation (see Implementation for full training parameters). After training, visual inspection of predicted masks revealed an accurate segmentation of colonies from background, although some errors remained around plate edges (Fig. 1b). This was not unexpected, considering that the Ilastik-generated masks also often contained artefacts at the edges. In order to circumvent this problem, we applied post-processing on the predicted masks, which effectively removed artefacts. The vast majority of resulting cropped regions contained a single colony; however, a few regions still contained multiple small, overlapping colonies. To reduce possible bias that might result from counting multiple colonies as one, we filtered these out during the classification stage.

For the classification task, we fine-tuned a Resnet-34 architecture that was pre-trained on ImageNet (http://www.image-net.org/)^8,14^, also implemented in the fast.ai library^6^. We trained the network using 1476 images of individually cropped colonies, which were split into five manually-labeled classes: white, red, pink, variegating, and multiple colonies. Again, 20% of colony images were kept aside for validation. After training, we achieved a validation accuracy of 91.8% across the five classes (Fig. 1d). Further, aggregating the three classes of non-white colonies together (red, pink and variegating) yielded a much higher validation accuracy of 98.6%. This higher pooled accuracy was encouraging, considering our desired output of percentages of white and non-white colonies per plate. It also demonstrates that most classification errors occur within non-white classes rather than between white and non-white classes, an expected result given that the non-white classes often do not have clear distinctions and can be difficult to define even by eye.

While we were encouraged by our high validation accuracy, we wanted to test the pipeline’s performance against manual counting in a real-world, experimental context. To this end, we took data from two published experiments testing trans-generational inheritance of *ade6*^*+*^ silencing in *S. pombe*^15^. In these experiments, *ade6*^*+*^ silencing was first induced by expression of small interfering RNAs (siRNAs) that are complementary to the *ade6*^*+*^ gene in a *paf1-Q264Stop* nonsense-mutant background, leading to red colonies. Paf1 is a subunit of the Paf1 complex (Paf1C), which represses siRNA-induced heterochromatin formation in *S. pombe*^16^. In the presence of the *paf1-Q264Stop* allele, the silenced (red) phenotype was inherited through meiosis, even in the absence of the original siRNAs that have triggered *ade6*^*+*^ repression. This was not the case if the progeny inherited a *paf1*^*+*^ wild type allele, i.e. the red silencing phenotype was lost. However, these white *paf1*^*+*^ cells inherited a marked *ade6*^*+*^ epiallele (*ade6*^*si3*^), which reinstated silencing when cells became mutant for Paf1 again in subsequent generations^15^. The following experiments were performed to quantify different aspects of this phenomenon.

In the first experiment, *paf1-Q264Stop* cells that inherited the *ade6*^*si3*^ allele and re-established the red silencing phenotype were plated on limiting adenine indicator plates to quantify the maintenance of this re-established silencing through mitosis. Out of 59 plates derived from a red progenitor colony, our automated pipeline predicted a mean of 84.7% non-white colonies, indicating a high degree of maintenance of *ade6*^*+*^ silencing. 10 of these plates were manually counted, and the mean difference in the percent of counted and predicted non-white colonies was 2.1 percentage points (Fig. 2a). As a control, colonies derived from cells of white *paf1-Q264Stop* progenitor colonies were also quantified by both methods. The automated pipeline predicted a mean of 0.57% non-white colonies across 60 plates, while manual counting of colonies on 12 plates detected 0 non-white colonies (Fig 2b).

**Figure 2.**
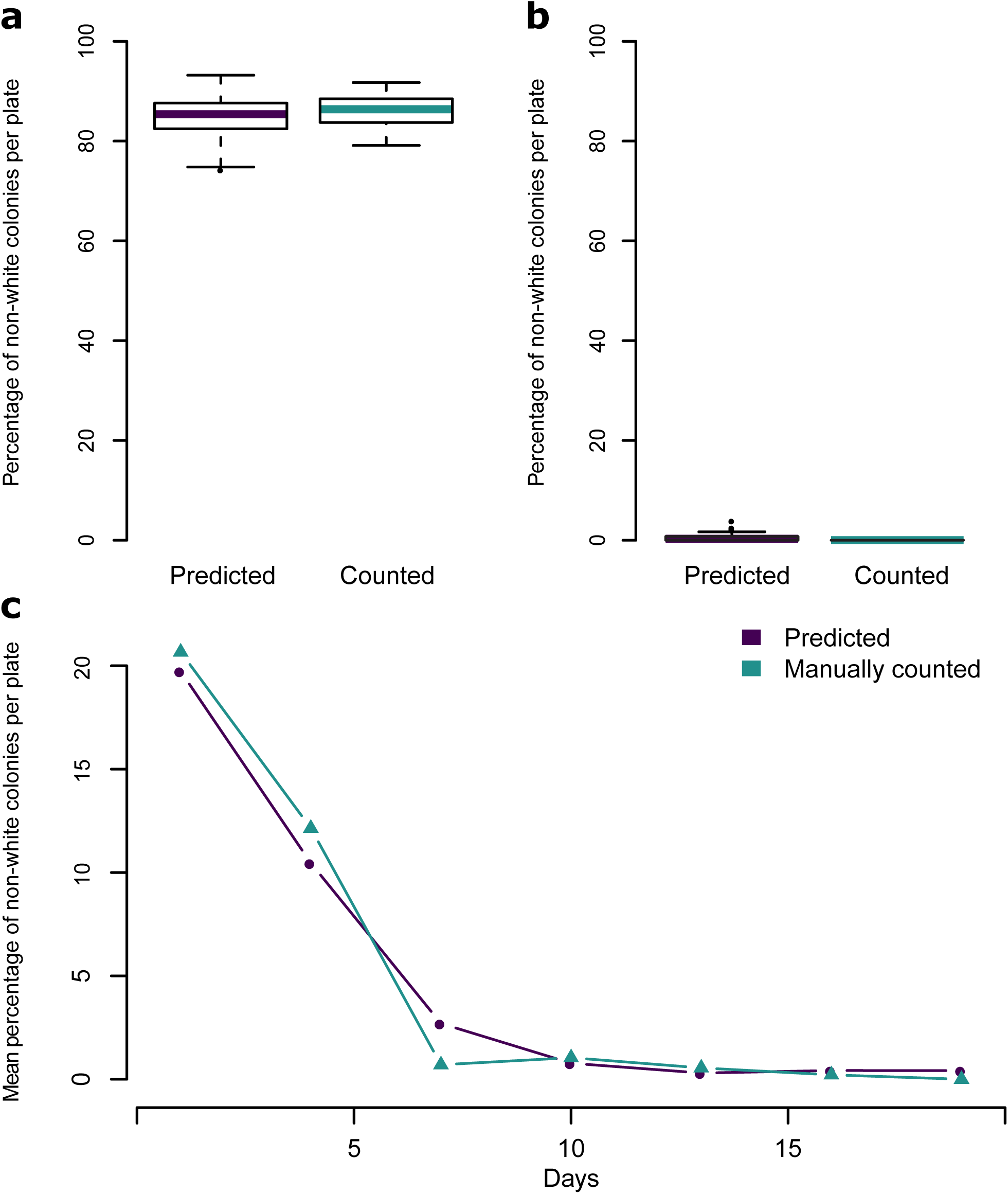
Comparison of automated colony classification and manual counting on experimental data. a) Boxplots showing the predicted or manually counted percentage of non-white colonies per plate on plates resulting from red *paf1-Q264Stop; ade6*^*si3*^ cells. n = 59 (predicted) or n = 10 (counted). b) Boxplots showing the predicted or manually counted percentage of non-white colonies per plate on plates resulting from white *paf1-Q264Stop* cells. n = 59 (predicted) or n = 12 (counted). For boxplots, center line is median, bottom and top hinges are 1^st^ and 3^rd^ quartiles, whiskers show the most extreme points within 1.5 times the interquartile range, and more extreme values are plotted as individual points. c) Mean predicted or manually counted percentages of non-white colonies per plate across a time course of mitotically dividing white *paf1*^*+*^ *ade6*^*si3*^ cells crossed to white *paf1-Q264Stop* cells every 3 days. The time course was repeated with 11 independent biological replicates, and 6 plates per replicate per timepoint were quantified by automated prediction (n=66 per time point). The mean of all 66 plates is reported for each timepoint. For manual counting, one plate per replicate per timepoint was counted (n=11 per time point); the mean of 11 plates is reported for each timepoint.

The second experiment was performed to assess mitotic stability of the *ade6*^*si3*^ epiallele. White *paf1*^*+*^ cells with an *ade6*^*si3*^ epiallele were grown exponentially and a sample was crossed to white *paf1-Q264Stop* cells every 3 days (30-40 mitotic divisions) over 19 days total. Progeny of these crosses were also observed to re-establish silencing; however, the frequency of re-establishment declined with the number of mitotic divisions the cells had gone through. The percentage of non-white colonies on plates resulting from crosses at each time point was both counted manually (66000 colonies total) and predicted using the automated pipeline. Both methods showed a near-exponential decrease in non-white colonies over time; the mean difference in the percent of non-white colonies between the two methods ranged from 0.26 to 5.1 percentage points (Fig 2c).

## Discussion and Conclusions

We observed several factors that contributed to the accuracy of the automated pipeline predictions versus manual counting. In general, the pipeline performed with very high accuracy on plates with a low (< 5%) percentage of non-white colonies; however, on some plates that had been grown for over two weeks, white colonies formed an irregular ring-like morphology and tended to be mis-classified as pink. The accuracy of the pipeline was also decreased for plates with very dense, small colonies. This may be partly due to the difficulty in segmenting individual colonies when they are touching one another, leading to many colonies being excluded from the analysis. Very small colonies are also more likely to be mistakenly filtered out during post-processing. Since cells with a silenced *ade6*^*+*^ gene tend to grow more slowly, this may lead to bias, as very small colonies are more likely to be non-white. However, plating cells at a controlled density and imaging the plates after an appropriate amount of time can counteract this potential bias. We have posted a full suggested protocol describing plating and imaging for most accurate prediction on Protocol Exchange (https://protocolexchange.researchsquare.com/.)

Our fully automated pipeline can be run on a CUDA-enabled CPU or GPU. On a GeForce GTX 1080 GPU, it took approximately 45 minutes to process 245 plate images, representing an 80-100x speedup over manual counting by an experienced biologist. In addition to saving time and manual labor, the pipeline has the potential to increase reproducibility by removing variations between individual researchers or computer monitors. Our pragmatic approach, combining transfer learning with readily available annotations, allows us to achieve accuracy comparable to human performance with relatively little training data. Our work should thus enable larger-scale experiments and higher statistical power, unlocking a true quantitative use of the red/white color assay in yeast research. We have made the code and trained network weights freely available to the community at https://github.com/fmi-basel/buehler-colonyclassification.

## ABBREVIATIONS

SGDR: stochastic gradient descent with restarts
SGD: stochastic gradient descent
MRI: magnetic resonance imaging
*S. pombe*: *Schizosaccharomyces pombe*
Paf1C: polymerase associated factor 1 complex
siRNA: small interfering RNA
*ade6*^*si3*^: *ade6+* epiallele marked with histone H3 lysine 9 tri-methylation and siRNAs
NCCR: national centers of competence in research
FMI: Friedrich Miescher institute for biomedical research
MTA: material transfer agreement
UBMTA: uniform biological material transfer agreement

## DECLARATIONS

### Availability of data and materials

Project name: Colony classification

Project home page: https://github.com/fmi-basel/buehler-colonyclassification

Operating system(s): Platform independent

Programming language: Python

Other requirements: Python 3.6 or higher, fastai library v. 0.7

License: GNU GPL v3.0

Any restrictions to use by non-academics: None

Raw images of plates with yeast colonies used in Figure 2 are available from the corresponding author on reasonable request.

### Author contributions

SHC trained and tested neural networks, wrote and optimized all scripts, compared manual vs. automated colony counting, and created figures. SHC and MB wrote the manuscript. LD optimized image acquisition and performed training and segmentation using Ilastik and Matlab, generating the training datasets. YS manually classified colony images. LD and YS gave feedback to help optimize the pipeline output.

## Acknowledgements

We thank M. Rempfler for discussions and for suggesting ideas for the automated segmentation, and K. Volkmann and S. van Eeden for help with image acquisition and processing.

## REFERENCES

1. Liu, W. et al. SSD: Single Shot MultiBox Detector. ArXiv151202325 Cs 9905, 21–37 (2016).

2. Lin, T.-Y., Goyal, P., Girshick, R., He, K. & Dollár, P. Focal Loss for Dense Object Detection. ArXiv170802002 Cs (2017).

3. Huang, J. et al. Speed/accuracy trade-offs for modern convolutional object detectors. ArXiv161110012 Cs (2016).

4. Hughes, A. J. et al. Quanti.us: a tool for rapid, flexible, crowd-based annotation of images. Nat. Methods 15, 587–590 (2018).

5. Sommer, C., Straehle, C., Köthe, U. & Hamprecht, F. A. Ilastik: Interactive learning and segmentation toolkit. in 2011 IEEE International Symposium on Biomedical Imaging: From Nano to Macro 230–233 (2011). doi:10.1109/ISBI.2011.5872394

6. Howard, J. & others. fastai. (GitHub, 2018).

7. van der Walt, S. et al. scikit-image: image processing in Python. PeerJ 2, e453 (2014).

8. He, K., Zhang, X., Ren, S. & Sun, J. Deep Residual Learning for Image Recognition. ArXiv151203385 Cs (2015).

9. Ronneberger, O., Fischer, P. & Brox, T. U-Net: Convolutional Networks for Biomedical Image Segmentation. ArXiv150504597 Cs (2015).

10. Huang, G. et al. Snapshot Ensembles: Train 1, get M for free. (2017).

11. Smith, L. N. Cyclical Learning Rates for Training Neural Networks. (2015).

12. Kumar, V. et al. Automated and real-time segmentation of suspicious breast masses using convolutional neural network. PLOS ONE 13, e0195816 (2018).

13. Dalmış, M. U. et al. Using deep learning to segment breast and fibroglandular tissue in MRI volumes. Med. Phys. 44, 533–546 (2017).

14. Deng, J. et al. ImageNet: A Large-Scale Hierarchical Image Database. IEEE Computer Vision and Pattern Recognition (CVPR) (2009)

15. Duempelmann, L. et al. Inheritance of a Phenotypically Neutral Epimutation Evokes Gene Silencing in Later Generations. Molecular Cell 74, 3 (2019).

16. Kowalik, K. M. et al. The Paf1 complex represses small-RNA-mediated epigenetic gene silencing. Nature (2015). doi:10.1038/nature14337

